# Stratified computational meta-analysis of 2213 acute myeloid leukemia patients reveals age- and sex-dependent gene expression signatures

**DOI:** 10.1101/494948

**Authors:** Raeuf Roushangar, George I. Mias

## Abstract

In 2018 alone, an estimated 20,000 new acute myeloid leukemia (AML) patients were diagnosed, in the United States, and over 10,000 of them are expected to die from the disease. AML is primarily diagnosed among the elderly (median 68 years old at diagnosis). Prognoses have significantly improved for younger patients, but in patients older than 60 years old as much as 70% of patients will die within a year of diagnosis. In this study, we conducted stratified computational meta-analysis of 2,213 acute myeloid leukemia patients compared to 548 healthy individuals, using curated publicly available data. We carried out analysis of variance of normalized batch corrected data, including considerations for disease, age, tissue and sex. We identified 974 differentially expressed probe sets and 4 significant pathways associated with AML. Additionally, we identified 70 sex- and 375 age-related probe set expression signatures relevant to AML. Finally, we used a machine learning model (KNN model) to classify AML patients compared to healthy individuals with 90+% achieved accuracy. Overall our findings provide a new reanalysis of public datasets, that enabled the identification of potential new gene sets relevant to AML that can potentially be used in future experiments and possible stratified disease diagnostics.

## INTRODUCTION

Acute myeloid leukemia (AML) is a heterogeneous malignant disease of the hematopoietic system myeloid cell lineage. AML is best characterized by the terminal differentiation in normal blood cells and excessive production and release of cells at various stages of incomplete maturation (leukemia cells). As a result of this faster than normal and uncontrolled growth of leukemia cells, healthy myeloid precursors involved in hematopoiesis are suppressed, and ultimately, can soar to death within months from diagnosis if untreated^1,2^. AML accounts for 70% of myeloid leukemia and nearly 80% of acute leukemia cases, making it the most common form of both myeloid and acute leukemia^2,3^. The number of new AML cases is increasing each year – in 2018 alone, there have been an estimated about 20,000 new diagnosed AML patients, over 10,000 of them will die from the disease^4^.

According to the 2016 World Health Organization (WHO) newly revised myeloid neoplasms and acute leukemia classification system^5^, AML prognosis criteria for classification is highly dependent on the presence of chromosomal abnormalities, including chromosomal deletions, duplications, translocations, inversions, and gene fusion. Mostly, AML is diagnosed through microscopic, cytogenetics, and molecular genetic analyses of patients’ blood and/or bone marrow samples. Microscopic examination is used to detect distinctive features (e.g. Auer rods) in cell morphology, cytogenetic analysis to identify chromosomal structural aberrations (e.g., t(8;21), inv(16), t(16;16), or t(9;11)), and molecular genetic analysis to identify gene fusion (e.g., RUNX1-RUNX1T1 and CBFB-MYH11), and mutations in genes frequently mutated in AML (e.g., NPM1, CEBPA, RUNX1, FLT3)^6-8^. These cytogenetic and molecular genetic analyses are used to identify prognosis markers that can be used to classify AML patients into three risk categories: favorable, intermediate, and unfavorable. The largest group of AML patients (almost 50%) however, present normal karyotype and lack genetic abnormalities^7-10^. These patients are classified as intermediate risk, and often have heterogeneous clinical outcome with standard therapy with risk of AML relapse^11^.

Additionally, AML prognosis worsens as age increases, and older patients respond less to current treatments with poorer clinical outcomes than their younger counterparts^12,13^. AML can occur in people of all ages but is primarily diagnosed among the elderly (>60 years), with a median age of 68 year at diagnosis^4^. Recent advances in AML biology expanded our understanding of its complex genetic landscape and led to significant improvement in prognoses and therapeutic strategy for younger patients^13,14^. However, in patients older than 60 years old, prognoses remain grim and therapeutic strategy has been nearly the same for more than 30 years^2,6,13-15^. Approximately 70% of AML patients 65 years of age or older die within a year following diagnosis^16^. While it is apparent that the nature of AML changes with age, still little is known about the extent of these associations and how they vary with patient’s age^14,17,18^. Taking into consideration age considerations in the identification of changes in AML global gene expression can lead to improved early diagnosis and improvement in treatment approaches for elderly patients. Further complicating, AML has multiple driver mutations and competing clones that evolve over time, making it a very dynamic disease^19,20^

Multiple gene expression analyses of AML have been carried out, 25 of which these have been systematically compared by Miller and Stamatoyannopoulos^21^, who analyzed information on 4918 genes, and identified 25 genes reported across multiple, with potential prognostic features. In this study, we performed comprehensive gene expression meta-analysis of 2213 acute myeloid leukemia patients and 548 healthy subjects using 34 publicly available gene expression microarray datasets (following strict inclusion criteria) to identify disease, sex- and age-related gene expression changes associated with AML. We identified sex- and age-related gene expression signatures that show similar alteration in gene expression levels and associated signaling pathways in AML and have used our results (gene sets) to predict AML or healthy status. We believe that our results may lead to improved AML early detection and diagnostic testing with target genes, which collectively can potentially serve as sex- and age-dependent biomarkers for AML prognosis compared to healthy, as well as the identification of new treatment targets with mechanisms of action different from those used in conventional chemotherapy

## RESULTS

### Data curation and gene expression preprocessing

We searched the Gene Expression Omnibus (GEO) public repository, based on our systematic workflow and inclusion criteria, Fig. 1a-b. Overall, 2,132 datasets were screened, 643 selected (577 were excluded as non-Affymetrix, various platform arrays). From the 66 remaining, 34 studies were excluded due to lack of metadata, non-peripheral blood and non-bone marrow tissues, cell line or cell-type specific, treated subjects). After this curation we obtained 34 age-annotated gene expression datasets from 32 different studies covering 2,213 AML patients and 548 healthy individuals. The sets were re-analyzed, starting from raw data, to perform a gene expression analysis of variance and functional pathway enrichment analysis (see online Methods). Table 1 provides a description on each dataset with a sub-table summary of all curated data used in our current study. After pre-processing each individual data set separately, Fig. 1b, we performed the statistical analysis on 44,754 probe sets which were common across all samples (Affymetrix expression array data).

**Figure 1.**
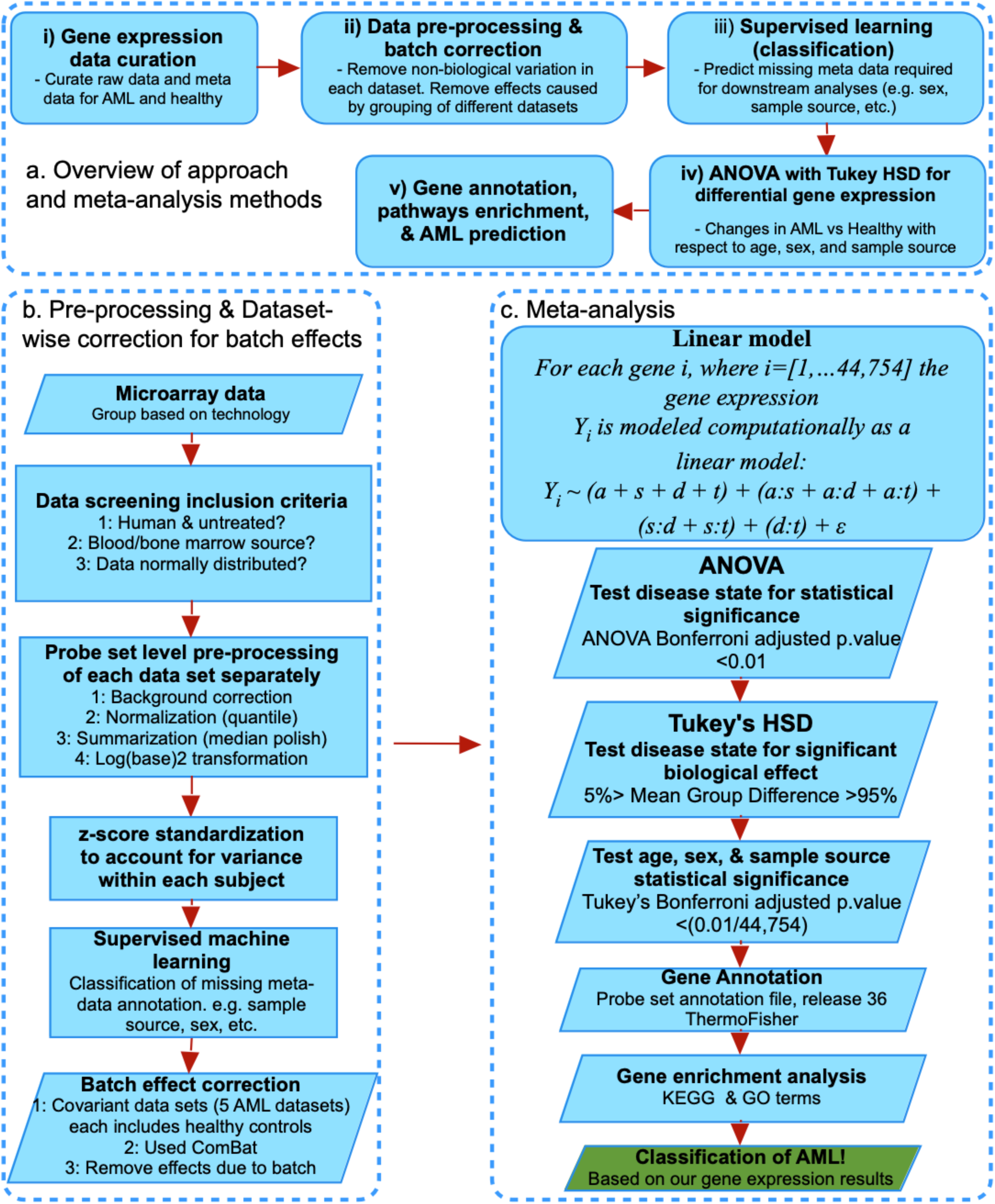
General approach, data curation, and analysis workflow summary. The flowchart shows in **(a)** the five main steps that summarize our method of approach for our study, and in (**b)** the curation and screening criteria for raw gene expression and annotation data files curation, data pre-processing, supervised machine learning for missing metadata prediction, and batch effects correction. (**c)** The meta-analysis included a linear model analysis of variance (ANOVA) coupled Tukey’s Honestly Significant Difference (HSD) post-hoc tests, and KEGG pathway and GO enrichment. Finally, we performed a machine learning classification of AML based on our findings.

**Table 1:**
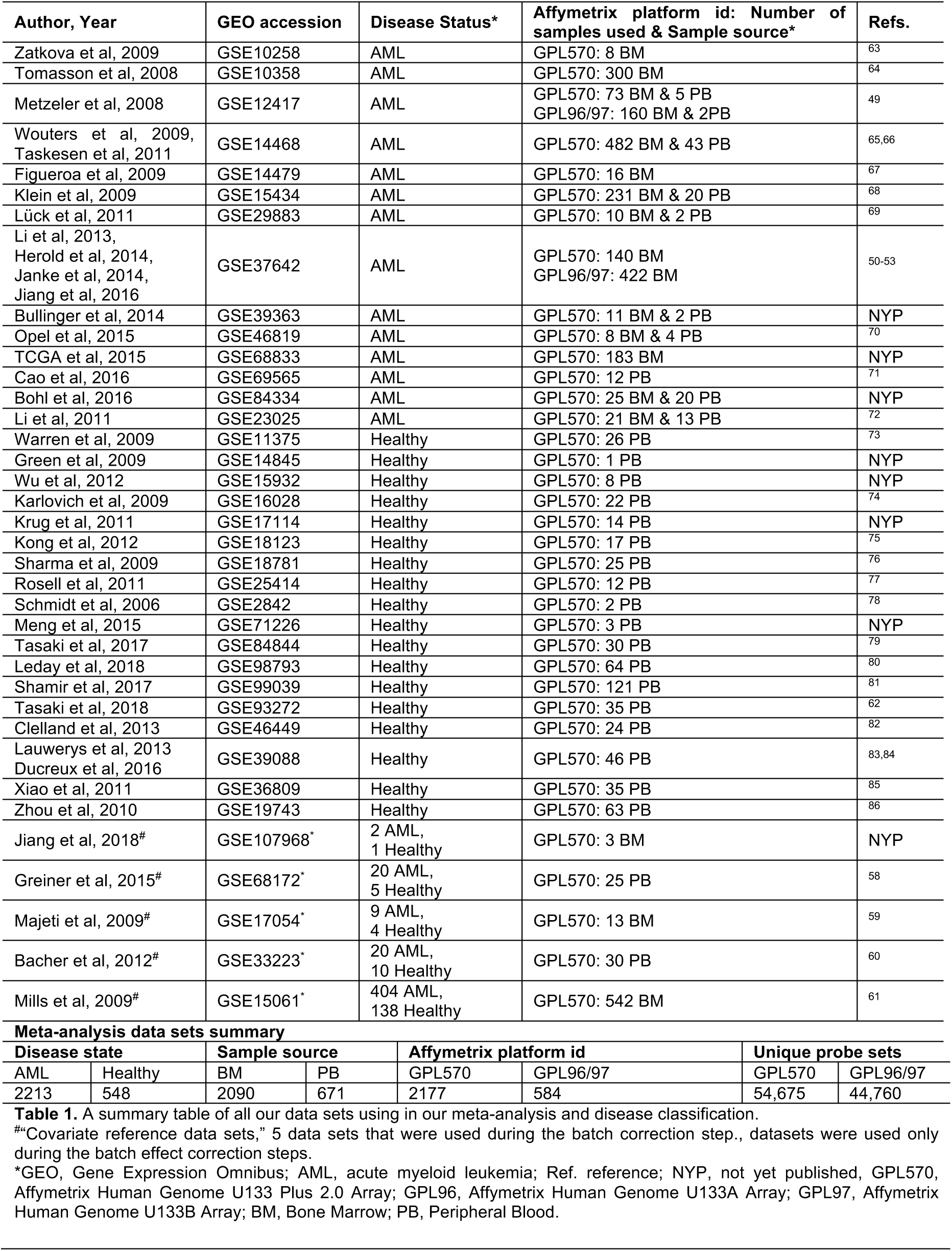
Summary table of all 34 gene expression datasets used in this study.

### Classification of missing metadata annotation

Following the data curation step, 805 arrays (802 AML and 3 healthy) of 2,761 curated data were found to be missing sex annotation, and 737 arrays (all AML patients) were missing information regarding the sample source (i.e. tissue, either bone marrow [BM] or peripheral blood [PB] annotation). To predict the missing sex and sample source meta-data, we trained and validated various machine learning supervised models, including logistic regression (LR) classification models. The prediction of missing annotations for these arrays was essential in our study, to increase the sample size, and statistical power^22^. The models were trained and verified using our annotated preprocessed expression data. Model training, parameters used in training, validation for this analysis are discussed in the Methods. Results from model training and predictions, including confusion matrix, model accuracy, and error can be viewed in Supplementary Table S1 online and results from classification for missing annotation are presented in Supplementary files 1 and 2 for sample source and sex annotations respectively.

### Batch correction

Our pre-processed data, AML and healthy, were processed using a “dataset-wise” batch effect correction approach. The datasets used in this study did not include within-study healthy controls, which would limit analysis of variance, and particularly the ability to separate biological from batch effects. To address this, we implemented an iterative batch effect correction approach, essentially employing a weight-based method for correcting batch effects. Assuming the batch effects due to each data set is a function of the number of samples in the data set (weight), normalizing sets of unevenly sized datasets may lead to unbalanced batch correction. We used 5 additional datasets as a reference set, which we refer to as “covariate” hereafter. Each of the covariate reference datasets included within study healthy controls. All 5 datasets together consisted of a total 613 arrays (455 AML and 158 healthy) (Table 2), and pre-processed exactly as our curated data sets. These were used together with each of the remaining datasets to batch correct each dataset with respect the covariate reference using ComBat^23^. After this dataset-wise correction, the 5 covariate reference datasets were removed, and our expression data were clustered using principal component analysis (PCA) to visually examine the effect of covariate reference datasets on distributing the batch weight during batch correction. The batch effect correction results were then compared to clustering results prior to batch effect correction (Supplementary Fig. 1)

**Table 2.**
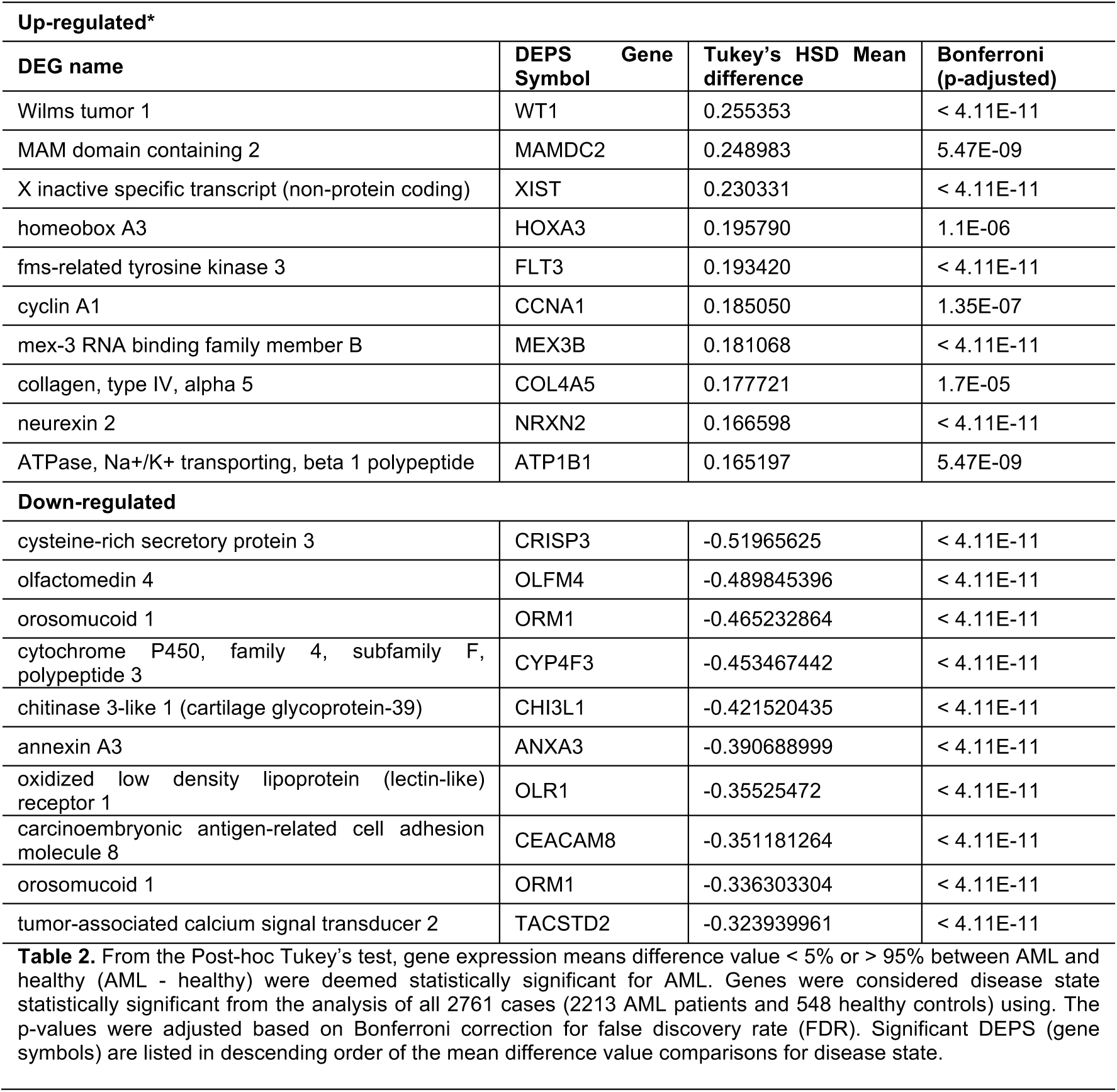
Top 10 up- and down-regulated of DEPS in AML from disease state.

### Analysis 1: Gene expression meta-analysis and enrichment analysis of AML disease state compared to healthy individuals

#### Gene expression meta-analysis of AML disease state

Following batch correction, we performed an analysis of differential expression (DE) on 34 data sets including 2213 AML patients and 548 healthy controls. Analysis of Variance (ANOVA)^24-26^ was performed according to a linear model (see method section **Meta-analysis**), including factors for age, sample source (adjust for differences in tissue between AML and healthy), and sex, as well as binary interactions thereof. In the analysis we used probe sets to avoid assumptions on averaging over multiple probe sets corresponding the same gene symbol. 974 Statistically significant differentially expressed probe sets (DEPS) (with genes corresponding to 964 unique gene symbols) for AML versus healthy were selected based on a Bonferroni^27^ adjusted p-value < 0.01 (accounting for multiple hypothesis testing), in conjunction with a two-tailed 5% quantile selection^28^ based on the mean difference distribution between AML-healthy group comparisons (post-hoc analyses using Tukey’s Honestly Significant Difference (HSD). The heatmap (Fig. 2a) shows the hierarchical clustering of genee expression from the 974 DEPS, including 487 up- and 487 down-regulated with respect to AML as compared to healthy. From this analysis, WT1 (Wilms tumor 1) with mean difference of 0.26 and adjusted p-value < 4.11×10^−11^ was the most DE up-regulated gene while CRISP3 (cysteine-rich secretory protein 3) with mean difference of −0.52 and adjusted p-value < 4.11×10^−11^ was the least DE gene. Figure 2b shows the top 10 up- and down-regulated DEPS with corresponding gene symbols, that resulted from this analysis (also listed in Table 2, including mean difference and Bonferroni p-adjusted values from post-hoc analysis using Tukey’s HSD tests). The entire list of all 974 DEPS can be found as Supplementary Table S2 online.

**Figure 2:**
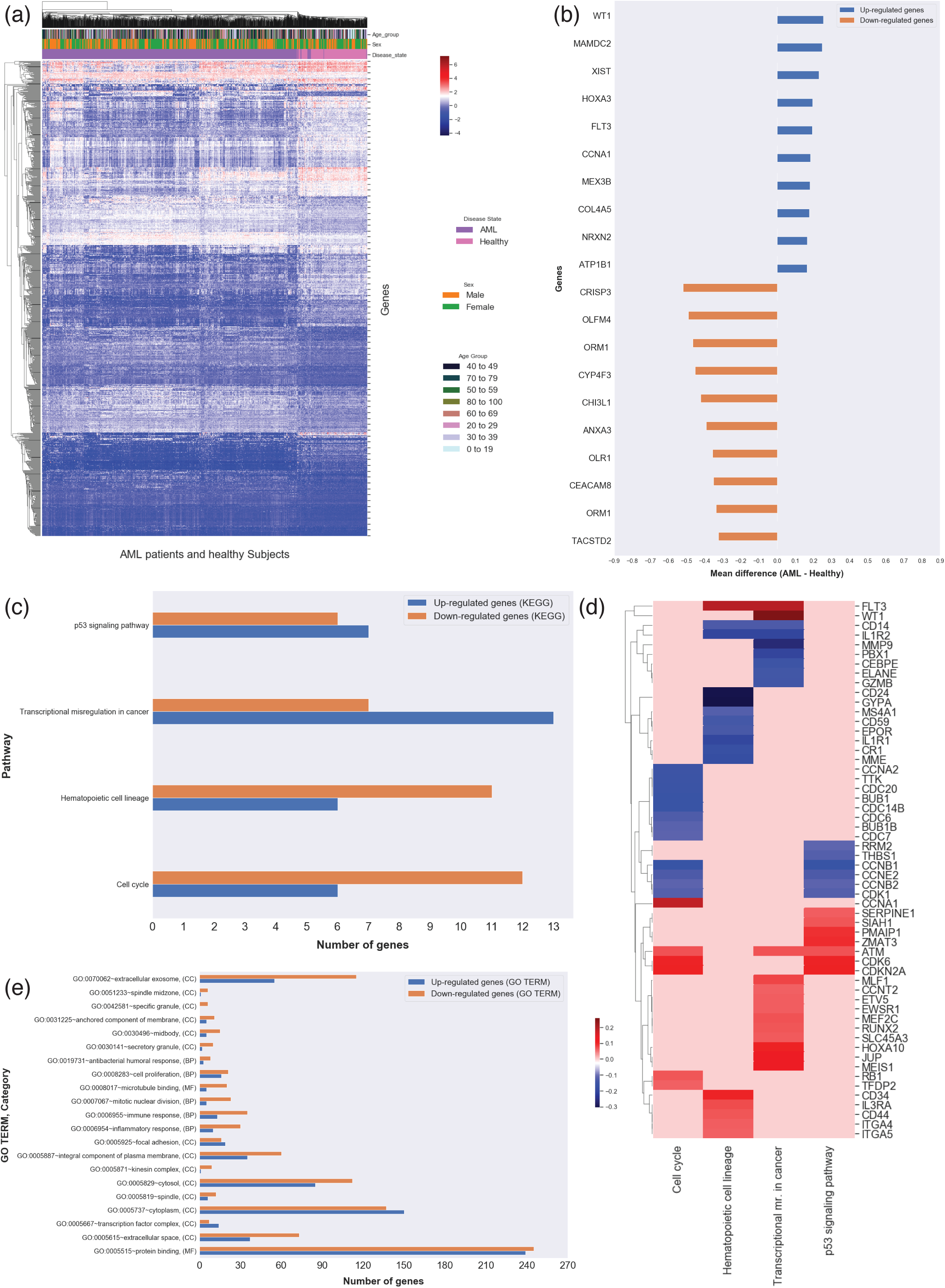
Functional classification of DEPS from AML meta-analysis and associated KEGG and GO enrichment analysis. For all panels, normalized values are represented in with blue for down-regulation and red for up-regulation, while light red/gray represents no reported specific direction. (**a)** Heatmap of 974 DEPS (rows) on 2,761 arrays (columns) including 2213 AML patients and 548 healthy individuals from AML meta-analysis, using unsupervised hierarchical clustering and Euclidean distance for clustering. The age of each individual is displayed at the bottom and illustrated in the color bar on the top (dark green for young and yellow for old). The disease state (AML vs healthy), sex of each subject and age-groups are represented in color bars on the top. (**b)** Horizontal barplot of the top 10 DEPS (gene symbols on vertical axis) from AML meta-analysis with mean difference values between AML and healthy (horizontal axis). Enrichment analysis identified 4 KEGG signaling pathways **(c)** for our AML DEPS, also visualized as a heatmap **(d)** of DEPS mean difference values between AML and healthy DEPS (rows) identified in these 4 KEGG signaling pathways (columns). The GO enrichment analysis results are summarized in **(e)**.

#### (ii)Gene enrichment analysis AML disease state DEPS

To identify signaling pathways associated DEPS in AML, gene enrichment analysis was performed on all 974 DEPS combined. Pathway over-representation analysis in Kyoto Encyclopedia of Genes and Genomes (KEGG)^29-31^ signaling pathways, and Gene Ontology (GO) term^32,33^ were carried out using the Database for Annotation, Visualization and Integrated Discovery (DAVID)^34,35^. Four KEGG signaling pathways were identified as enriched (Benjamini and Hochberg^36^ adjusted p-value < 0.05), including Hematopoietic cell lineage, Cell cycle, p53 signaling pathway, and Transcriptional misregulation in cancer. The 4 KEGG signaling pathways are summarized in Table 3 (see also Supplementary Fig. 2a-d), including unadjusted p-values and Benjamini and Hochberg^36^ adjusted p-values. 56 DEPS including 27 up- and 29 down-regulated (Fig. 2c) were associated these signaling pathways, and the heatmap of their mean differences is shown in Fig. 2d. From our gene enrichment analysis for overrepresented biological GO terms, 21 GO terms were statistically significant with 727 DE unique identities (335 up- and 392 down-regulated). GO terms included protein and microtubule binding for the molecular function (MF) category, inflammatory and immune responses, mitotic nuclear division, and cell proliferation response for the biological process (BP) category, and finally, cytoplasm, extracellular exosome, cytosol, extracellular space, integral component of plasma membrane immune response, and others, for the cellular component (CC) category (Fig. 2e). The entire list of our enrichment analysis results (statistically significant over-representation in KEGG and GO terms) can be found as Supplementary Table S3 online.

**Table 3.**
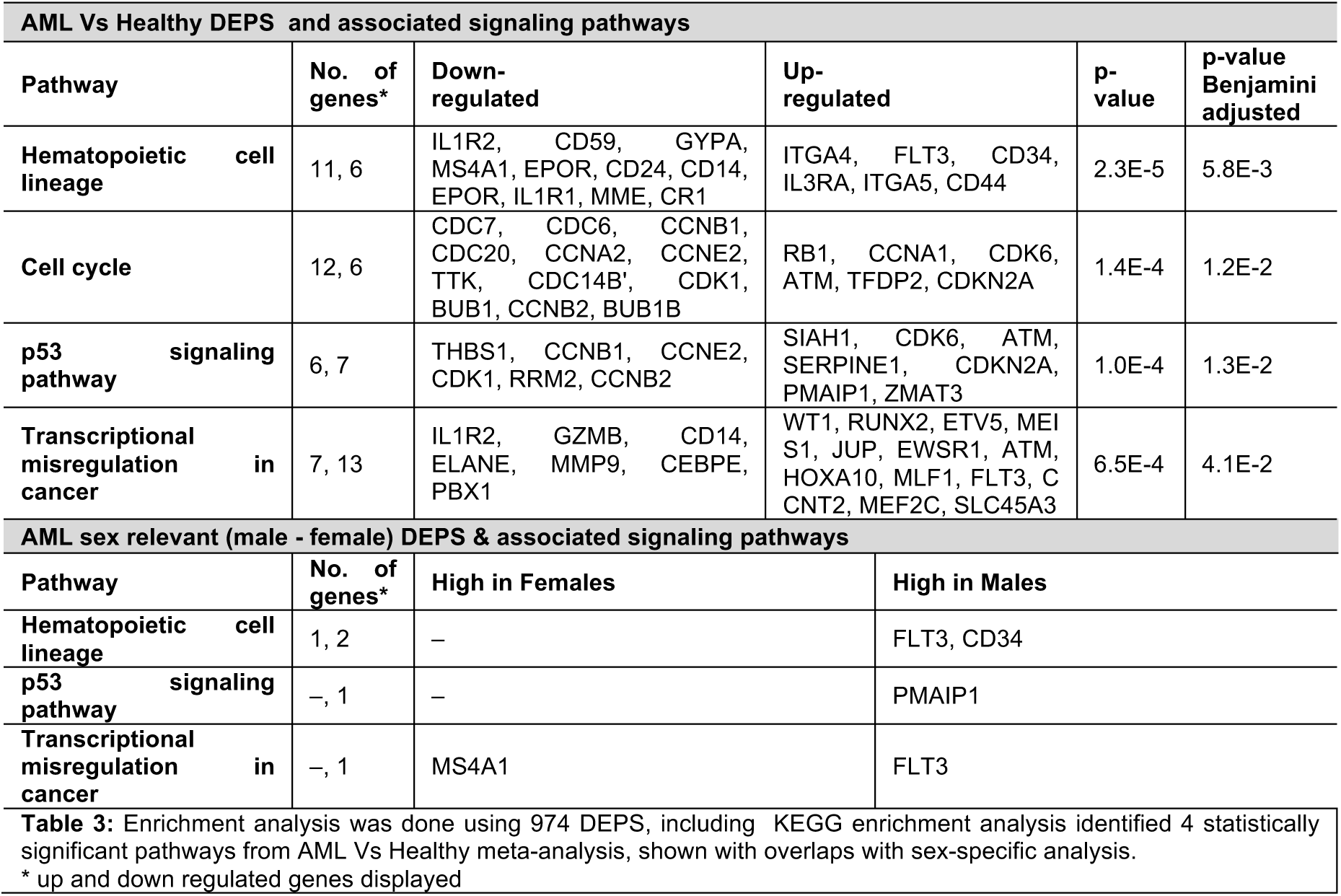
KEGG pathway analysis of DEPS from meta-analysis of 34 gene expression datasets.

### Analysis 2. Gene expression meta-analysis and enrichment analysis of sex- and age-related DEPS in AML

Further analysis of gene expression and pathways enrichment were conducted in order to characterize sex- and age-specific gene expression changes in AML patients compared to healthy individuals, (i) **Analysis 2a:** “**Sex-relevance differential gene expression meta-analysis and associated signaling pathways in AML”,** and (ii) **Analysis 2b:** “**Age-dependent differential gene expression meta-analysis and associated signaling pathways in AML”**. We used the same filtering criteria in both analyses as those used in analysis 1 for significant DEPS and signaling pathways between AML patients and healthy controls. In addition, DEPS were regarded as statistically significantly (up- or down-regulated) for each factor, sex and age, if they displayed Bonferroni adjusted p-value from Tukey’s HSD < 2.2×10^−7^ (=0.01/44,754 probe sets tested).

#### Analysis 2a. Sex-relevance differential gene expression meta-analysis and associated signaling pathways in AML

Gene expression meta-analysis was also used to identify DEPS that show sex differences between male AML patients as compared to female AML patients. 266 DEPS were regarded statistically significant (p-value < 2.2×10^−7^). A list of all 266 DEPS (including whether higher in either males or females, gene title and symbol, male-female mean difference, and Bonferroni corrected p-value) can be found as Supplementary Table S3 online. 70 DEPS were found to overlap between analysis 1 (AML disease state) and analysis 2 (Sex-relevance in AML). Figure 3a shows these 70 DEPS with gene symbol annotations, and their mean difference values in the heatmap, which displays differences in significance for a common DEPS in both analyses 1 and 2. Figure 3b shows the hierarchical clustering of the 70 DEPS (rows) on sex and disease state of all 2,213 AML and 548 healthy subjects (columns) indicated by color bars above the heatmap. The top 10 DEPS higher in either males or females from this analysis are shown in Figure 3c.

**Figure 3:**
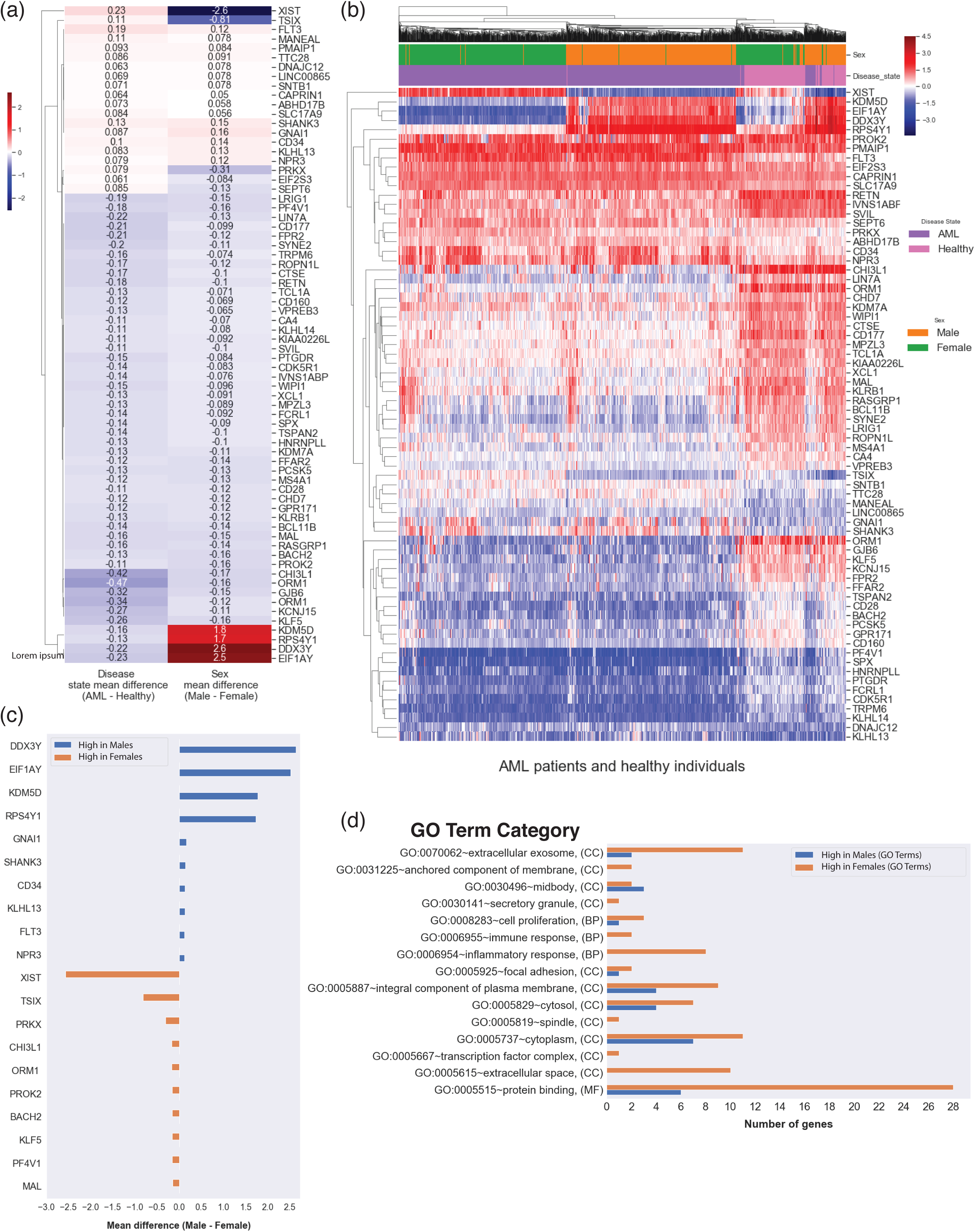
Sex-related gene expression meta-analysis in AML. **(a).** The heatmap of mean difference values comparison between the 70 DE overlapping genes between Analysis 1 and Analysis 2a. **(b)** Heatmap the 70 DEPS expression (rows) on 2,761 arrays (columns) including 2213 AML patients and 548 healthy individuals from Analysis 2a of sex-relevance in AML (using unsupervised hierarchical clustering and Euclidean distance for clustering). The disease state (AML vs healthy) and sex of each subject are indicated in color bars at the top. **(c).** Horizontal barplot of the top 10 DEPS (gene symbols on vertical axis), with the mean difference values between male-female (horizontal axis). (**d).** Enrichment analysis for statistically significant overrepresented biological GO terms on the 70 DE genes.

For enrichment analysis, we searched for common intersections in KEGG pathways and GO terms between the sex meta-analysis and the 974 DE probe sets from disease state in AML meta-analysis. Sex-relevant DEPS were found in 3 different signaling pathways, including, genes higher expressed in males FLT3 and CD34 in Hematopoietic cell lineage, FLT3 in Transcriptional misregulation in cancer 1, and PMAIP1 in p53 signaling pathway 1, and MS4A1 was higher in females and found in Hematopoietic cell lineage pathway (Table 3). Figure 3d shows GO analysis results, where 15 overrepresented biological GO terms were overlapped, including terms for extracellular space, immune response, protein binding, spindle, and midbody. The entire list of our enrichment analysis (statistically significant KEGG and GO terms) can be found as Supplementary Table S4.

#### Analysis 2b. Age-dependent differential gene expression meta-analysis and associated signaling pathways in AML

The subjects were binned in 8 age-groups: 0-19, 20-29, 30-39, 40-49, 50-59, 60-69, 70-79, and 80-100 years old. From this meta-analysis, 1395 unique probe sets across all age-groups were identified as statistically significant (Bonferroni adjusted p-value < 2.2×10^−7^) (Supplementary Table S5). From these 375 unique DEPS (372 unique gene symbols) were found to overlap with the 974 DEPS probe sets from our AML disease state meta-analysis, accounting for an overall 1400 binary comparisons between the multiple age groups deemed statistically significant, based on Tukey HSD tests between age-group pairs. The entire list of 1400 identified pairwise differences between age groups and associated probe set/gene information can be found as Supplementary Table S6 online. The top 10 up- and down-regulated DEPS (labeled with gene symbols) from this analysis are shown in Fig. 4a. Additionally, Fig. 4b shows 75 DEPS with gene symbols identified to have appeared specifically in one age-group comparison. Utilizing results for KEGG analysis for signaling pathways from analysis 1, Fig. 4c shows 17 DE genes identified in all 4 KEGG pathways according to age groups (see also Table 4).

**Figure 4:**
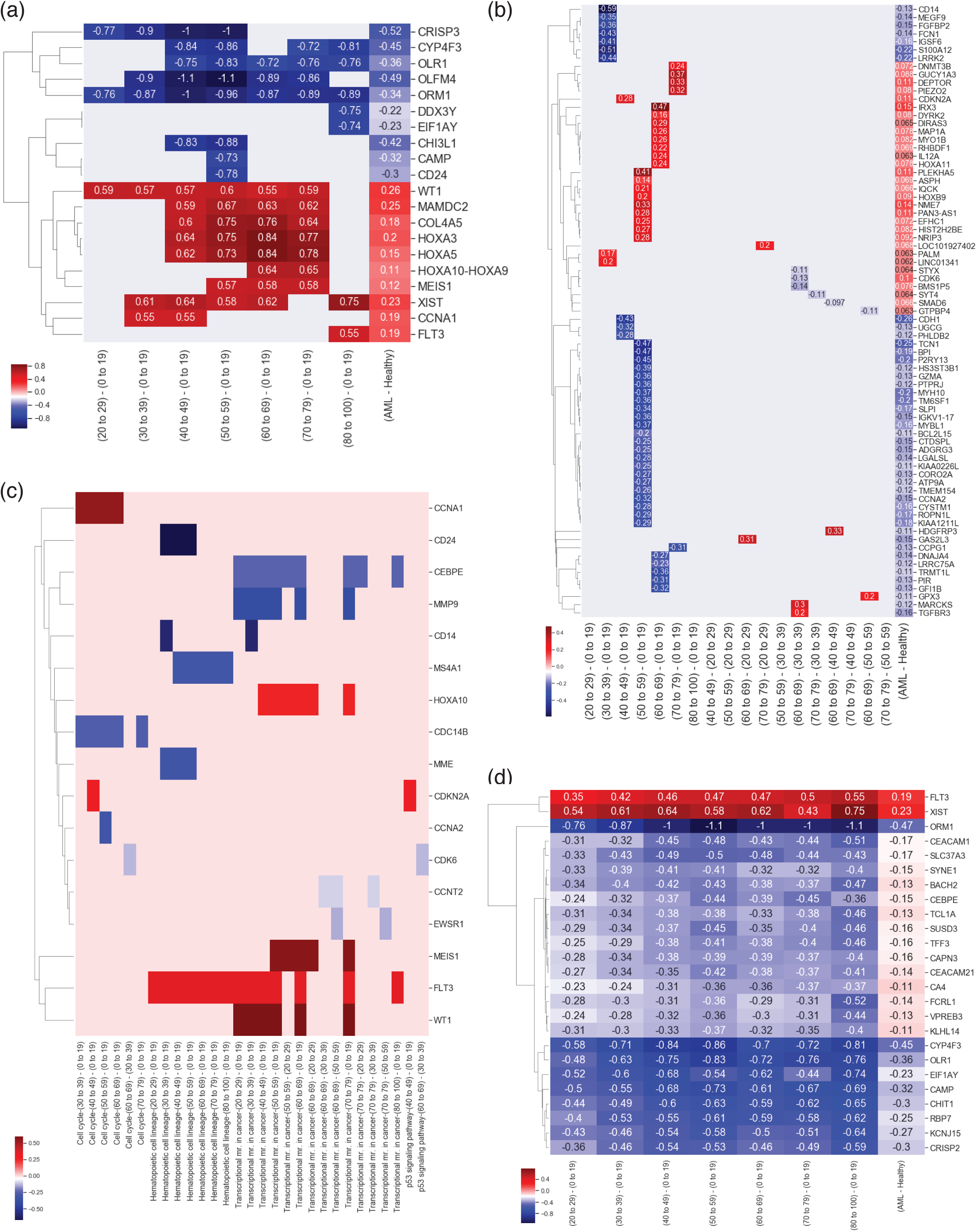
Age-related gene expression meta-analysis in AML. **(a)** The top 10 up- and down-regulated DEPS overlapping AML and age-related analyses. 75 DEPS specific to a single age-group comparison, **(b)**. **(c)** The mean difference of 25 DEPS with respect to the 0-19 baseline across all other groups are plotted to illustrate changes with aging. We note that the mean difference values between AML and healthy cohorts are shown in the right-most column of panes (**a)-(c)** for reference comparisons. (**d)** Overlaps over KEGG pathways of 17 DE genes identified in 4 KEGG pathways according to age groups.

**Table 4.**
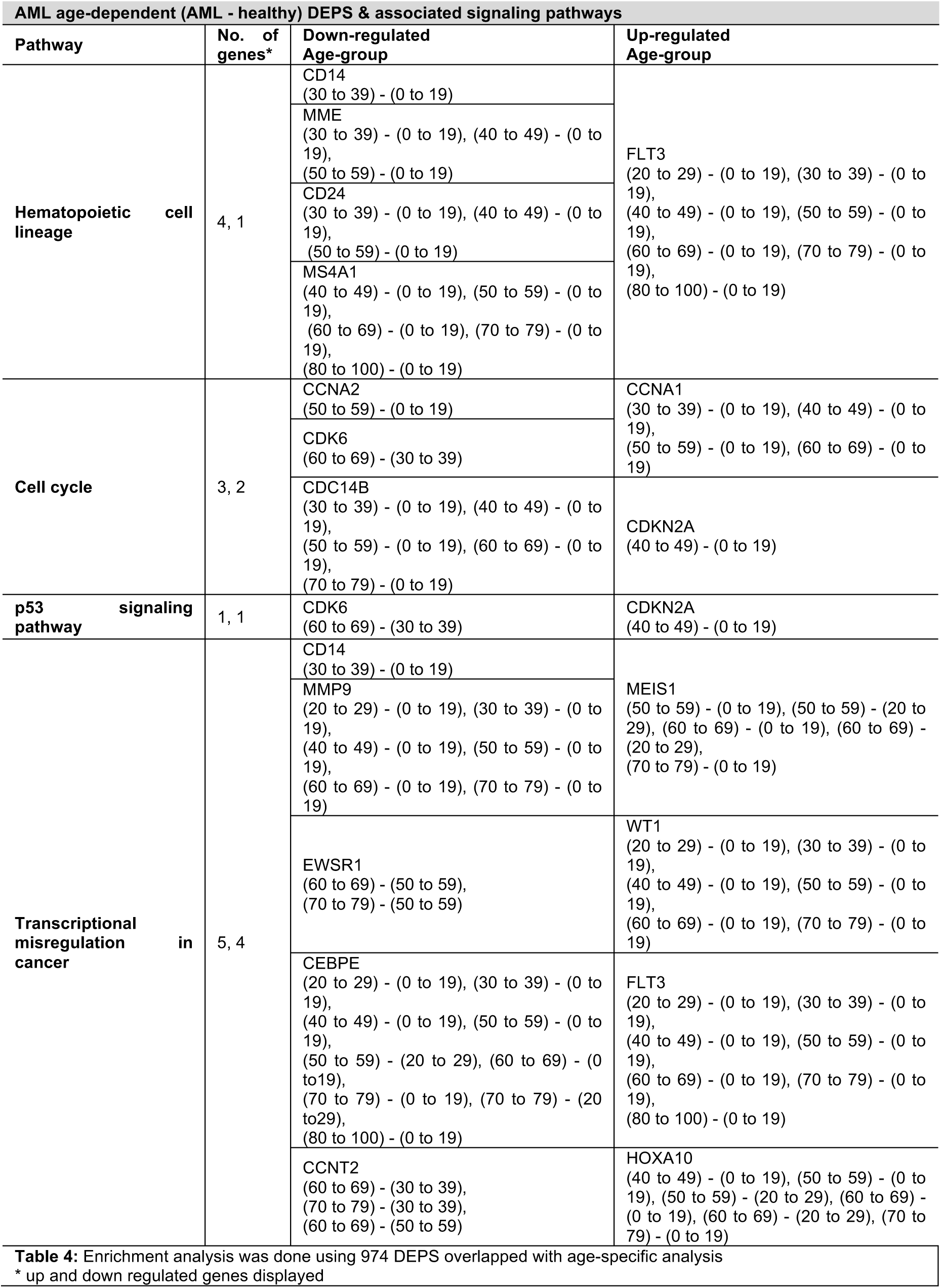
KEGG pathway analysis of DEPS from meta-analysis of 34 gene expression datasets overlap with age-specific findings.

To investigate further the progression with age, pairwise correlations between age-groups were computed. The 0-19 age-group was used as a common comparison reference with respect to other groups. Using this 0-19 group as a baseline, Figure 4d shows the mean difference of 25 DEPS with respect to the 0-19 baseline across all other groups. The mean difference values between AML and healthy are shown in the right-most column of Fig. 4a, b and d for reference.

### AML Classification Machine Learning Model

We used the 974 DEPS to train a k-nearest neighbor (KNN) algorithm in ClassificaIO^37^. All 34 datasets (16 AML and 18 healthy) were used for training, and testing was done on all 5 covariate reference datasets, include AML and healthy subjects. The KNN algorithm trained was 98% accurate, and >90% accurate in testing results (see online Methods for parameters and also details in Supplementary File 3).

## DISCUSSION

In the present study, we aimed to establish, disease sex-linked and age-dependent biomarkers from genes with similar changes in gene expression levels and associated signaling pathways relevant to AML. Utilizing microarray gene expression data and combined with various machine learning models, respectively, our biomarkers were indicative of prognostic signature for AML prediction compared to healthy with 90+% achieved accuracy. We re-analyzed data aggregated from our curation of 34 publicly available microarray gene expression datasets covering 2213 AML patients and 548 healthy individuals to identify changes in AML gene expression associated with disease state (AML compared to healthy), sex-linked (male compared to female), and age-dependent (across age-groups compared to baseline).

We performed 3 differential probe set (gene) expression and gene enrichment analyses, as discussed below. We note here that our study identified multiple potentially significant DEPS, with age and sex related differences associated with AML. While our findings may generate further hypothesis-driven investigations, we need to also identify the study’s limitations: primarily the analysis of AML and healthy subjects involved bone-marrow and blood samples respectively in each disease group. We tried to account for this utilizing tissue as an effect in our linear model, and including multiple interactions. Other limitations include an unbalanced AML/healthy ratio, as well as the lack of in-study healthy controls. To address these we attempted to account for batch effects using a dataset-wise iterative batch correction transformation, as discussed in the methods. Finally, we also included binary interactions between the factors in the analysis to account for interaction-related confounding effects.

*i) Analysis 1 Gene expression meta-analysis and associated signaling pathways of AML disease state compared to healthy individuals*, was carried out to identify DEPS in AML disease state. The results from this analysis were then used as baseline indicator for AML disease state. 974 DEPS (487 up- and 487 down-regulated) were identified as significantly differentially expressed between AML patients and healthy individuals (Bonferroni adjusted p-value < 0.01). Among these 6 genes are known to be involved in AML functional pathways, including 4 up-regulated, JUP, CCNA1, FLT3, PIK3R1, and 2 down-regulated, CD14, CEBPE. The top 10 up- and down-regulated genes from this analysis are listed in Table 2 with their respected Tukey’s HSD mean difference and Bonferroni p-adjusted values. As shown in Figure 2b of the top 10 up- and down-regulated DEPS and corresponding gene annotations -- WT1 (Wilms tumor 1) was found to be the most expressed and CRISP3 (cysteine-rich secretory protein 3) was the most under-expressed gene. WT1 is a transcriptional regulatory protein essential to cellular development and cell survival, and it has been known to be highly expressed with an oncogenic role in AML^38,39^, in agreement with our findings. However, CRISP3’s direct role in AML is still under investigation. CRISP3 is a member of the cysteine-rich secretory protein CRISP family with major role in female and male reproductive tract, and is mainly expressed in salivary gland and bone marrow^40^. Recently, 80 genes were reported as “extracellular matrix specific genes” in leukemia, and CRISP3 was among the downregulated DE genes reported^41^. CRISP3 associations with AML merit further investigation.

The enrichment analysis for GO terms of the 974 DE probe sets (Fig. 2c) results showed 727 identifiers (335 up- and 392 down-regulated) enriched for 21 GO terms. 592 of these (257 up- and 335 down-regulated) were enriched in the cellular component (CC) categories mainly associated with cytoplasm, extracellular exosome, cytosol, and extracellular space. These terms are rather generic, but may still reflect relevance to AML development and progression^42,43^. Biological process (BP) category, GO terms included inflammatory and immune responses, and cell proliferation, which are expected as AML is characterized by terminal differentiation of normal blood cells and excessive proliferation and release of abnormally differentiated myeloid cells, and likely affects many biological processes associated to the immune system. The four statistically significant KEGG pathways identified in the pathway enrichment analysis encompassed 56 DEPS (Table 3). Transcriptional misregulation in cancer was the most up-regulated pathway in AML (13 up-regulated DE genes, while Hematopoietic cell lineage, and Cell cycle pathways were mostly down-regulated, and the p53 signaling pathway was balanced in terms of up/downregulated DE genes (Fig. 2c). The enriched pathways Fig. 2d shows the mean difference values of the 56 DE pathway-associated genes, including 27 genes up- and 29 down-regulated. These KEGG pathways are known to be involved in tumorigenesis. Additionally, the majority of the associated DE genes from AML meta-analysis with the identified signaling pathways are known to be abnormally expressed in AML. These findings are consistent with findings from other studies and our current understanding of AML pathogenesis.

The DEPS overlap with the 25 genes reported by Miller and Stamatoyannopoulos that were reported in at least 8 studies^21^, namely HOXA10, CD34, MEIS1,VCAN, RBPMS and MN1. In terms of the genes reported in the same study for poor progression we also consistently identified as upregulated HOXA10, RBPMS, CD34, GNAI1, CLIP2, DAPK1, GUCY1A3, ANGPT1 and FLT3, and as downregulated UGCG. While these are known markers, with consistent expression differences, our additional results need to be investigated further and experimentally validated, including mechanistic considerations.

*ii) Analysis 2a Sex-dependent gene expression meta-analysis and associated signaling pathways in AML compared to healthy individuals*, was performed to explore the relevance of patients’ sex on gene expression and to identify sex-linked genes and associated signaling pathways in AML. A total of 266 DEPS were found statistically significant in this analysis, with 70 found to overlap with the DEPS from Analysis 1 (Fig 3a-b). The top10 up- and down-regulated DE genes with respect to females include (Fig. 3c) – DDX3Y (DEAD-Box Helicase 3 Y-Linked), EIF1AY (Eukaryotic Translation Initiation Factor 1A Y-Linked), KDM5D (Lysine Demethylase 5D), RPS4Y1 (Ribosomal Protein S4 Y-Linked 1) with higher expression in males compared to females, and XIST (X Inactive Specific Transcript), TSIX (TSIX Transcript, XIST Antisense RNA), and PRKX (Protein Kinase X-Linked) were as higher in females. These genes are known to be sex-specific and show such differences and sex separation within the AML and the healthy groups respectively (Fig. 3d). The role of these genes as positive controls in studies with AML needs to be investigated further. We also reported sex and AML known genes that were statistically significant in our analysis, including FLT3 and MAL.

*iii) Analysis 2b Age-dependent gene expression meta-analysis and associated signaling pathways in AML compared to healthy individuals*, was carried out to identify common set of age-dependent genes and associated signaling pathways and to explore age-dependent trends in gene expression in AML. The age-dependent meta-analysis in AML using ANOVA, identified 1,395 DEPS (Bonferroni adjusted p-value <0.01). To identify age-related DEPS in AML we overlapped the 1,395 DEPS to our findings of 974 DEPS in AML disease state (Analysis 1) (Fig. 4a), and identified an overlap of 375 DEPS (Bonferroni adjusted p.value <0.01). As shown in Figure 4b, the top 10 most and least DE age-associate genes in AML according to the mean difference values in seven age-groups, including their corresponding values from AML disease state in column “AML - healthy” for comparisons. Interestingly, CRISP3 was among the down regulated genes specifically and involved in this analysis as well, specifically associated with differences in younger age groups, 20 to 49 years of age as compared to 0 to 19 age group. Other genes showing age-specific differences included HOXA3, HOXA5 and HOXA10-HOXA9, which belong to the homeobox genes (HOX) family of transcription factors, essential to embryonic development and hematopoiesis, and associated with chromosomal abnormalities translocation and over-expression in AML^44,45^. Also identified with age-specific DE, was ORM1, which in Analysis 1 was among the top-10 most under-expressed genes, and was also among the 70 DE genes in analysis 2a. ORM1’s direct role in AML also merits further investigation, given ORM1 involvement in immunosuppression and inflammation^46^. Finally, we have identified 75 DEPS that show association with only one age-group, exclusively from all other age-groups, suggestive of potential age-specific differential gene expression signature.

In summary, our study successfully integrated multiple datasets to perform a study of gene expression in AML, across multiple factors that included disease, sex and age considerations, and identified interesting genes, both known and not previously reported as differentially expressed in each factor. We identified 974 DEPS and 4 associated significant pathways involved in AML, and 70 sex- and 375 age-related DE signatures. Using the 974 DEPS, a KNN model allowed AML with 91.7% accuracy. We hope that these findings may provide additional relevant targets for further experimental mechanistic studies, and to help identify new markers and therapeutic targets for AML.

## METHODS

The generalized workflow consisted of five main steps: i) Curation of microarray gene expression data, ii) Preprocessing of raw data files followed by batch effect correction, iii) Predictions of missing annotation data using supervised machine learning, iv) Differential gene expression analysis, and v) Gene enrichment for pathway analysis that includes gene annotation, and finally gene expression-based prediction of AML (Fig. 1a).

### Gene expression data curation and screening criteria

Datasets used in this study were selected from the GEO public repository, maintained by the National Center for Biotechnology Information (NCBI)^47^ (https://www.ncbi.nlm.nih.gov/geo/). To facilitate speed of search and keep up-to-date with possible new and relevant datasets, as soon as they were released, a Python script was used that utilized functions from the Entrez Utilities from Biopython^48^. We used the script to navigate the GEO records, and download microarray gene expression datasets up to 10/18. We additionally utilized Python packages, including Pandas, NumPy, and Matplotlib for data structure, numerical computing for data processing, and data visualization respectively. We used strict inclusion criteria to maintain consistency in each dataset selection, screen for availability of both raw and meta-data annotation files provided, human samples used from untreated subjects, and that the sample source was from either bone marrow (BM) and/or peripheral blood (PB). Array platform was restricted to Affymetrix, which was found to have the most available data, and to avoid cross-platform normalization issues. Inclusion criteria and the data curation workflow are illustrated in Fig. 1 a-b.

### Gene expression data sets used in our analysis

The curation method is summarized in the Supplementary File 4 flowchart and in the Results section. For our analysis we included 34 age-dependent datasets from 32 different studies, 16 included AML and 18 healthy subjects respectively. From the 34 datasets, 32 were produced from Affymetrix GeneChip Human Genome U133 Plus 2.0 (GPL570) and 2 conducted on Affymetrix GeneChip Human Genome U133 Array Set (GPL96 & GPL97) arrays. Table 1 provides detailed information about each data set, including the number of samples used from each dataset, sample tissue source, as well as the total number of AML patients and healthy subjects. Two studies, GSE12417^49^ and GSE37642^50-53^, were originally conducted on two different Affymetrix array types (GPL570, and GPL96 & GPL97), so each was separated into two subgroups and each subgroup was considered as individual dataset in our meta-analysis, data set GSE12417: (i) subgroup 1 included 73 BM and 5 PB samples, and (ii) subgroup 2 included 160 BM and 2PB. For dataset GSE37642 (i) subgroup 1 included 140 BM and (ii) subgroup 2 422 BM samples (Table 1).

### Dataset annotation and preprocessing

Figure 1b outlines the workflow of our preliminary data analysis including preprocessing. For each dataset used in our analysis, raw microarray CEL files were downloaded from GEO, metadata was reviewed, and the data was manually curated to guarantee that and each array, which corresponded to either an AML patient or healthy individual, was verified and correctly annotated for sample source (BM or PB), platform technology used, age, sex, and disease state (AML or healthy). Raw CEL files from individual datasets were individually pre-processed using the RMA (Robust Multi-Array Average) algorithm^54-56^. Datasets with mixed sample source, i.e both BM and PB, were pre-processed together irrespective of sample source. Preprocessing consisted of correction for background noise using RMA background correction on perfect match (PM) raw intensities, quantile normalization to obtain the same empirical distribution of intensities for each array, median polish summarization of probes into probe sets to estimate gene-level expression value, and logarithm base-2 transformations of gene expression values to facilitate data interpretation (normal distributions) and comparisons between arrays. Additionally, our expression data were first reduced to 44,754 probe sets that are common to and appeared in all data. Data sets were z-score standardized across all probe sets and arrays.

### Prediction of missing sex- and sample source annotations from curated data sets

805 arrays (802 from AML patients and 3 were healthy subjects) of curated data were not annotated for sex, while 737 arrays (all AML patients) were missing sample source information. Without these metadata, we would have to discard the data, which in turn would limit the statistical power for the study, and our ability to correct for biases stemming from individual datasets^22^. To address this, we used supervised machine learning classifiers to predict metadata. For all prediction, we used ClassificaIO^37^, a machine learning for classification user interface, which we recently developed, to carry out the machine learning classification analyses utilizing the sklearn package in Python^57^

To predict sex pre-processed data sets, 1956 arrays (including both healthy and AML), that include 44,754 probe sets and their annotated sex information were used to train logistic regression (LR) classification models, and to predict 805 sex annotations. Additionally, 2024 arrays were used to train for sample source, and the prediction was performed on 737 arrays.

The supervised machine learning LR classifier we used with the following parameters:

> random_state = None, shuffle = True, penalty = l2, multi_class = ovr, solver = liblinear, max_iter= 100, tol = 0.0001, intercept_scaling = 1.0, verbose = 0, n_jobs = 1, C = 1.0, fit_intercept = True, dual = False, warm_start = False, class_weight = None

The trained models for classification of missing sex and sample source annotation from curated data achieved > 95% classification accuracy with ~ 3-5% classification errors. Confusion matrix details, model accuracy and error for training and testing are presented in Supplementary Table S1 online, and results in Supplementary files 1 and 2. To account for training overfitting, we used 10-fold cross-validation on all 1,956 gene expression data arrays for training and validation.

### Dataset-wise correction approach for batch effects correction

Batch correction was done using a dataset-wise correction. Here we refer to the term “dataset-wise correction,” to indicate performing batch correction iteratively on one dataset at a time, against a reference set of datasets chosen to account for variability. We used this approach to account for the lack within-study healthy controls in the curated gene expression datasets. To address this issue, we used 5 additional datasets the included within-study controls, GEO accessions: GSE107968, GSE68172^58^, GSE17054^59^, GSE33223^60^, and GSE15061^61^(Table 1B). We refer to the latter datasets hereafter as “covariate” reference datasets, as they were as the reference datasets in the batch correction. Our approach aimed to balance/distribute the weight of batch effects exerted by each dataset, as this is dependent on the number of observations within a given dataset. Combined, the covariate reference datasets included 613 total arrays, totaling 455 AML and 158 healthy controls. We used ComBat^23^ to correct for study batch effects, as its empirical Bayes-based algorithm uses both scale and mean center based methods, providing an appropriate algorithm^23^. Covariate reference datasets were treated as the covariate for batch during batch correction, to improve performance in correcting for batch effects rather than biological variation. After batch correction, we used principal component analysis (PCA), visualizing components in both 2 and 3 dimensions, to compare the clustering results for corrections. Covariate reference datasets were removed after the batch correction step and were not part of our downstream meta-analysis. (Supplementary Fig. S1).

### Gene expression meta-analysis

After batch correction step, we performed gene expression meta-analysis for differential expression on the merged datasets (34 data sets, 16 AML and 18 healthy), where the expression values for all 44,754 common probe sets were aggregated. The effects of patients’ age, sex, and sample source, including their pairwise interactions were investigated using an analysis of variance (ANOVA)^8,62^. For each gene *i*, where *i*=[1,…44,754], the gene expression probe set *Y*_*i*_ was modeled computationally as a linear model:

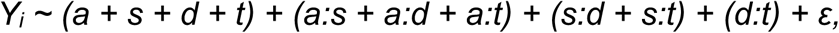

where *d* is the disease state (AML or healthy), *a* is age (between 0 to 100 years), *s* is sex (female or male), *t* is sample source (BM or PB), and *ε* is a random error term. We note that the model includes sample source and its interactions to address comparisons involving different tissues in AML and healthy subjects (BM or PB respectively).

From the ANOVA analysis, genes were deemed to be disease state statistically significant (differentially expressed) if they displayed ANOVA Bonferroni-adjusted p-value < 0.01. Post-hoc analysis for significant genes was conducted for comparisons (between groups) using Tukey’s Honestly Significant Difference (HSD) tests. Additionally, we performed a quantile-based effect filter, were genes were deemed to show biological effects in our analysis if they displayed mean difference values in the <5% and/or > 95% quantiles of the mean difference distributions of the binary group comparisons. Based on the post-hoc analysis, genes were deemed to be statistically significantly (up- or down-regulated) if they displayed Tukey HSD using a Bonferroni adjusted cutoff for p-value < 0.01/44,754.

### Functional and pathway enrichment analysis

We carried our enrichment analysis for DEPS using the Database DAVID^34,35^, the KEGG database^29-31^ for signaling pathways, GO terms functional annotation for over representation of biological function ^32,33^ were utilized and signaling pathways were deemed significant based on Benjamini-Hochberg adjusted p-value < 0.05.

### Using a k-nearest neighbor model to predict AML

Before gene expression data passed to the k-nearest neighbor (KNN) algorithm to train, gene expression signatures resulted from our meta-analysis were used to extract expression values. KNN in ClassificaIO^37^ was used to carry out this analysis. All 34 data sets (16 AML and 18 healthy) were used for training, and testing was done on all 5 covariate data sets, include AML and healthy subjects. Dependent, target, and testing data files were prepared in accordance with ClassificaIO^37^ user guide. The KNN model used the following parameters (Supplementary File 3):

> random_state = None, shuffle = True, metric = minkowski, weights = uniform, algorithm = auto, n_neighbors = 5, leaf_size = 30, n_jobs = 1, p = 2, metric_params = None

The trained model was 98% accurate, while testing was 91.7% accurate (details of training and testing are given in Supplementary File 3.

## Supporting information

## DATA AVAILABILITY STATEMENT

The datasets generated in the study, supplementary data, tables, figures and files are available online at http://doi.org/10.5281/zenodo.1492796

Datasets re-analyzed in the study are publicly available on the Gene Expression Omnibus repository, at https://www.ncbi.nlm.nih.gov/geo/ under accessions summarized in Table 1.

## ACKNOWLEDGEMENTS

R.R. has been supported by The Paul and Daisy Soros Fellowship for New Americans. G.I.M. and research reported in this publication have been supported by a Jean P. Schultz Endowed Biomedical Research Fund award, and by NIH/NHGRI Grant HG0006785. The content is solely the responsibility of the authors and does not necessarily represent the official views of the National Institutes of Health and other funders.

## AUTHOR CONTRIBUTIONS STATEMENT

R.R. and G.I.M. wrote the main manuscript text and prepared the figures. All authors reviewed the manuscript.

## ADDITIONAL INFORMATION

### Competing interests

G.I.M. has consulted for Colgate-Palmolive North America and received compensation. R.R. declares no potential conflict of interest.

