## Supplementary material for "Stratified computational meta-analysis of 2213 acute myeloid leukemia patients reveals age- and sex-dependent gene expression signatures"

### A. Supplementary Figures

#### Supplementary Figure S1: Principal component analysis.

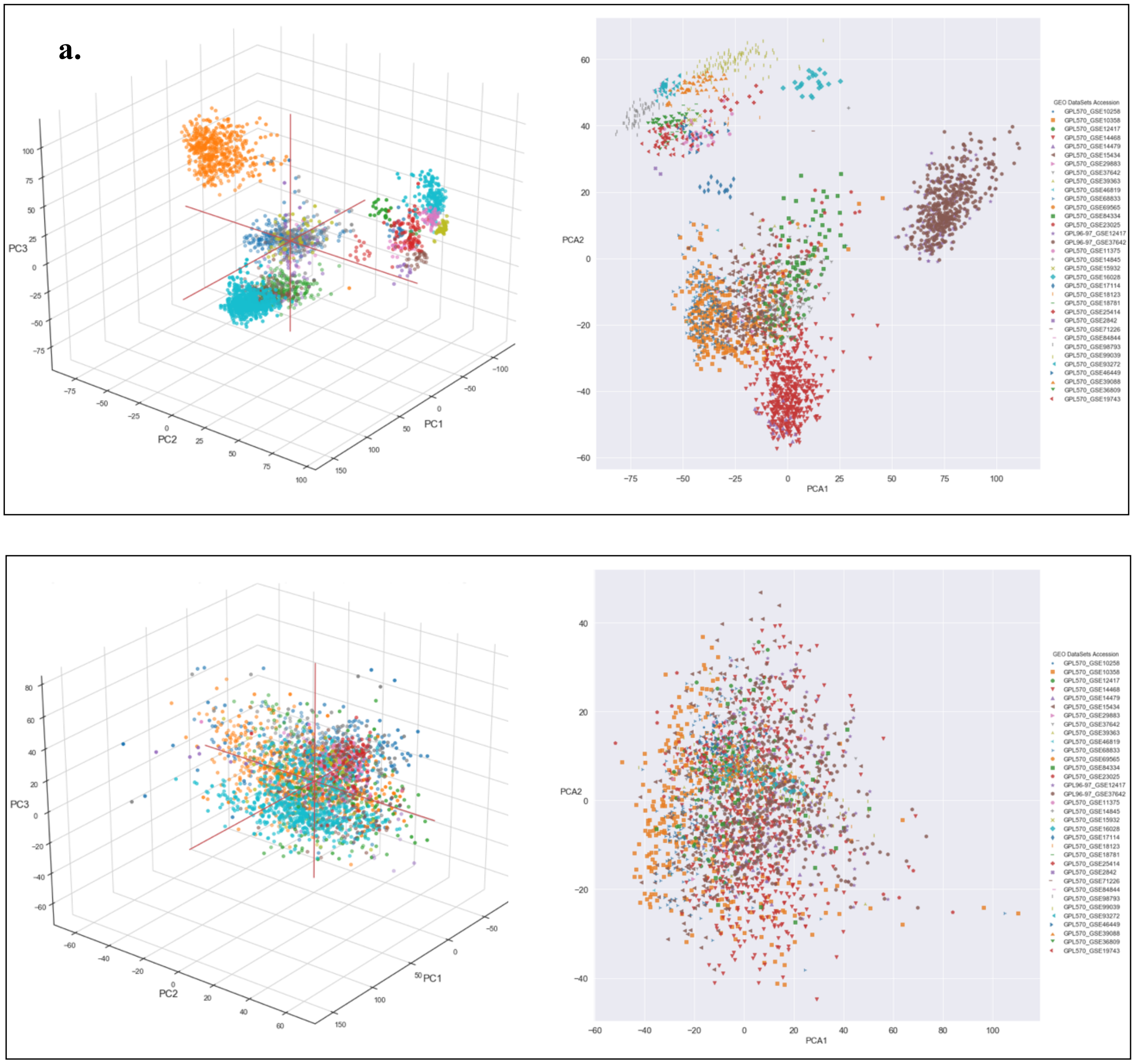

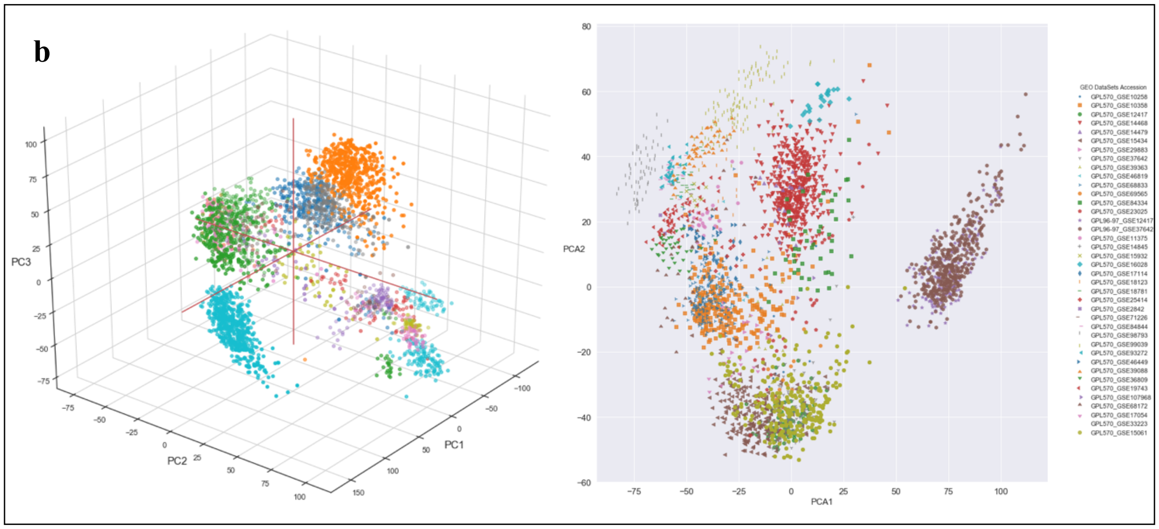


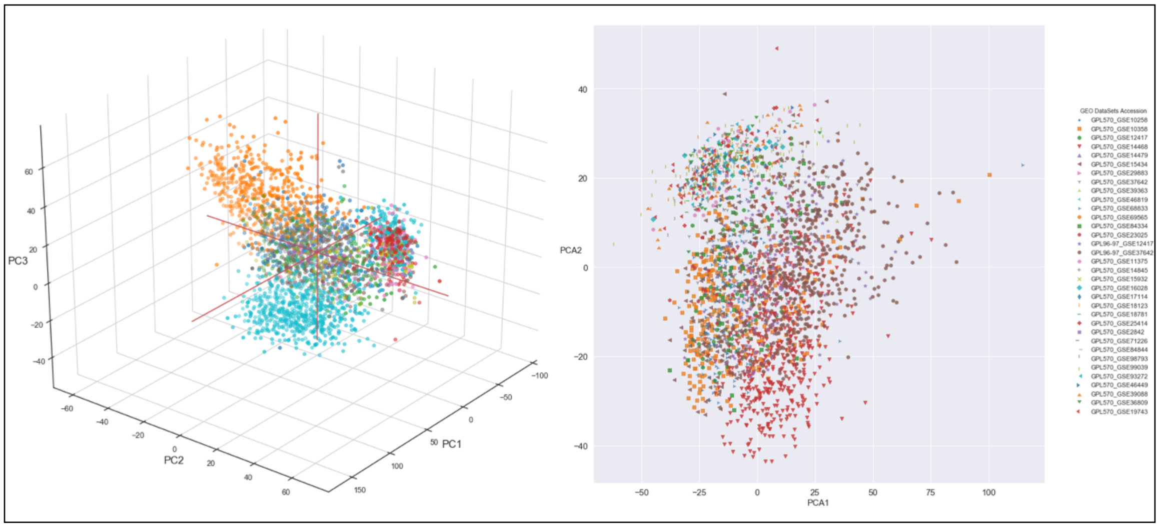


Principal Component Analysis (PCA) of pre-processed, logarithm base-2 transformed, z-score standardized data sets with 44,754 probe sets across all data sets: **(a)** 2761 subjects (AML and healthy), and data before batch correction **(top)**. The data was corrected without covariates resulting to loss of biological effect information due to lack of within-sample controls (**bottom**). **(b)** Data shown before batch effect correction including 2761 subjects (AML and healthy), and additionally 613 covariate datasets (5 data sets), (labeled green) **(Top)**, and, **(bottom)** PCA of, data post batch effect correction using ComBat as descripted in the methods section.

#### B. Supplementary Tables

Supplementary Tables are provided as sheets within the Supplementary_Tables.xlsx Excel File. These include:

1. ***Supplementary Table S1***. Logistic Regression Classification Model Parameters and Training Result Summary
2. ***Supplementary Table S2***. Analysis 1: Gene expression meta-analysis of AML disease state
3. ***Supplementary Table S4***. Analysis 2a. Sex-relevant differential gene expression meta-analysis and associated signaling pathways in AML
4. ***Supplementary Table S4***. Analysis 1: Gene enrichment analysis of AML disease state differentially expressed genes
5. ***Supplementary Table S5***. Analysis 2b. Age-dependent differential gene expression meta-analysis and associated signaling pathways in AML. 1389 DEG
6. ***Supplementary Table S6***. Analysis 2b. Age-dependent differential gene expression meta-analysis and associated signaling pathways in AML. 372

#### *C. Supplementary Files*

*The following supplementary files are provided:*

1. ***Supplementary_Tables.xlsx***: Supplementary Tables Spreadsheets
2. ***SupplementaryFile1_SampleSourceClassification.xlsx***: Logistic Regression Training and Testing Results for Sample Source Classification.
3. ***SupplementaryFile2_SexClassification.xlsx***: Logistic Regression Training and Testing Results for Sex Classification.
4. ***SupplementaryFile3_AMLClassification.xlsx***: KNN Model Training and Testing Results for AML Classification.

#### D. Online Data Availability

Supplementary data, tables, figures and files are available online at https://www.zenodo.org/badge/DOI/10.5281/zenodo.1492796.

| | File Name | Description |
|---|---|---|
| 1 | AML_and_Healthy_archive_with_predicted_sex_and_sample_source_for_ANOVA_with_age_groups_and_with_shorter_ID_REF_for_U133AB.csv | Information of 2,761 cases (2,213 AML and 548 healthy) in our analysis: age, sex, sample source, and disease state |
| 2 | All_613_AML_and_Healthy_with_44754_probsets_RMA_Normalized_Log2Trans_Zscore_Standardized_Transposed_Data.csv | Control data used for ComBat batch correction |
| 3 | All_2761_AML_and_Healthy_with_44754_probsets_RMA_Normalized_Log2Trans_Zscore_Standardized_Transposed_Data.csv | 2,761 normalized gene expression data of 44,754 probe sets |
| 4 | All_2761_Corrected_for_All_Factors_SampleSource_DiseaseState_Batch_Datasetwise_2213_AML_1st_548_Healthy_2nd_and_removed_613_dummy_with_44754_probsets_RMA_Normalized_Log2Trans_Zscore_Standardized_Transposed_Data_with_shorter_ID_REF_for_U133AB.csv | 2,761 batch corrected gene expression |
| 5 | ANOVA_P_Values_For_All_2761_Corrected_for_All_Factors_SampleSource_DiseaseState_Batch_Datasetwise_2213_AML_1st_548_Healthy_2nd_and_removed_613_dummy_with_44754_probsets_RMA_Normalized_Log2Trans_Zscore_Standardized_Transposed_Data.csv | ANOVA result for all 2,761subjects |
| 6 | Tukey_DiseaseState_P_Values_Corrected_for_All_Factors_SampleSource_DiseaseState_Batch_Datasetwise_multiplied_diff_by_1_corrected.csv | Tukey's Honest Significant Difference test result for AML vs healthy |
| 7 | Tukey_Sex_P_Values_Corrected_for_All_Factors_SampleSource_DiseaseState_Batch_Datasetwise.csv | Tukey's Honest Significant Difference test result for female vs male |
| 8 | Tukey_SampleSource_P_Values_Corrected_for_All_Factors_SampleSource_DiseaseState_Batch_Datasetwise.csv | Tukey's Honest Significant Difference test result for bone marrow vs normal blood |
| 9 | Tukey_Age_P_Values_Corrected_for_All_Factors_SampleSource_DiseaseState_Batch_Datasetwise_with_Probesets_as_Rows_Name.csv | Tukey's Honest Significant Difference test result for age groups |
| 10 | 1956_Dependent_Data_Subjects_with_Sex_Info_from_all_2761_all_2761_expression_arrays_after_BECorrection | Gene expression dependent data for sex classification (1956 data points) |
| 11 | 1956_Target_Data_For_Dependent_Data_Subjects_with_Sex_Info_from_all_2761_expression_arrays_after_BECorrection.csv | Gene expression target data for sex classification (1956 data points) |
| 12 | 805_Testing_Data_Subjects_with_No_Sex_Info_from_all_2761_expression_arrays_after_BECorrection.csv | Gene testing for sex classification (805 data points) |
| 13 | 2024_Dependent_Data_Subjects_with_SampleSource_Info_from_all_2761_expression_arrays_after_BECorrection.csv | Gene expression dependent data for sample source classification (2024 data points) |
| 14 | 2024_Target_Data_For_Dependent_Data_Subjects_with_SampleSource_Info_from_all_2761_expression_arrays_after_BECorrection.csv | Gene expression target data for sample source classification (2024 data points) |
| 15 | 737_Testing_Data_Subjects_with_No_SampleSource_Info_from_all_2761_expression_arrays_after_BECorrection | Gene testing for sample source classification (737data points) |
| 16 | GPL570_HG-U133_Plus_2.txt | Gene/probe set conversion annotation file for Affymetrix GPL570 array |
| | ***SUPPLEMENTARY FILES*** | |
| 17 | ***SupplementaryInformation*** | Manuscript Supplementary Information |
| 18 | ***Supplementary_Tables.xlsx*** | Supplementary Tables Spreadsheets |
| 19 | ***SupplementaryFile1_SampleSourceClassification.xlsx*** | Logistic Regression Training and Testing Results for Sample Source Classification. |
| 20 | ***SupplementaryFile2_SexClassification.xlsx*** | Logistic Regression Training and Testing Results for Sex Classification. |
| 21 | ***SupplementaryFile3_AMLClassification.xlsx*** | KNN Model Training and Testing Results for AML Classification. |
