## Supplementary material for "Stratified computational meta-analysis of 2213 acute myeloid leukemia patients reveals age- and sex-dependent gene expression signatures"

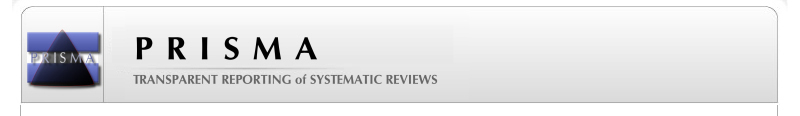
**PRISMA 2009 Flow Diagram**

**Screening**

**Included**

**Eligibility**

**Identification**

Records identified through database searching (GEO)
(n = 2,132 datasets)

Additional records identified through other sources
(n = 0 )

Records after duplicates removed
(n = 0 datasets )

Records screened
(n=643 datasets)

Records excluded

Non Affymetrix/various array platforms
(n = 577 datasets)

Full-text articles assessed for eligibility
(n = 66 datasets)

Full-text articles excluded,

(treated/lacking age information/non peripheral blood or bone marrow/cell type specific)
(n = 34 datasets)

Studies included in qualitative synthesis
(n = 32 studies)

Studies included in quantitative synthesis (meta-analysis)
(n = 32 studies)
